# Learning to be on time: temporal coordination of neural dynamics by activity-dependent myelination

**DOI:** 10.1101/2021.08.17.456520

**Authors:** Afroditi Talidou, Paul W. Frankland, Donald Mabbott, Jérémie Lefebvre

## Abstract

Activity-dependent myelination is the mechanism by which myelin changes as a function of neural activity, and plays a fundamental role in brain plasticity. Mediated by structural changes in glia, activity-dependent myelination regulates axonal conduction velocity. It remains unclear how neural activity impacts myelination to orchestrate the timing of neural signaling. We developed a model of spiking neurons enhanced with neuron-glia feedback. Inspired by experimental data and use-dependent synaptic plasticity, we introduced a learning rule, called the Activity-Dependent Myelination (ADM) rule, by which conduction velocity scales with firing rates. We found that the ADM rule implements a homeostatic control mechanism that promotes and preserves synchronization. ADM-mediated plasticity was found to optimize synchrony by compensating for variability in axonal lengths by scaling conduction velocity in an axon-specific way. This property was maintained even when the network structure is altered. We further explored how external stimuli interact with the ADM rule to trigger bidirectional and reversible changes in conduction delays. These results highlight the role played by activity-dependent myelination in synchronous neural communication and brain plasticity.

## 1 Introduction

Neural communication is supported by oligodendrocytes, a type of glial cell that produces a white lipid-rich substance called myelin. Like a conductor, myelin wraps around axonal membranes, influencing the conduction velocity of action potentials. Axonal conduction velocity is a critical feature of the nervous system because it determines conduction delays (i.e., the time it takes for an action potential to reach its destination). The distribution of such delays promotes a rich repertoire of dynamics [1–4]. While often overlooked, axonal conduction delays play a critical role in neural communication due to the integrative and synchronous nature of neural signaling. Synchrony is defined as coincident neural activity (i.e., temporal alignment of action potentials) and represents an important challenge for neural circuits, especially in presence of axons of various lengths. This is distinct from the concept of oscillatory synchrony in which periodicity is also present. Multiple converging action potentials travel along different paths, yet still manage to arrive synchronously at their destination. Synchrony is thus crucial not only for neuronal communication (e.g., post-synaptic integration), but also for synaptic plasticity. Inadequate or incomplete myelination (e.g., in white matter injury) may lead to jittered action potential timing and conduction lags resulting in suboptimal integration in postsynaptic cells and thus impacting the whole downstream neural signaling [5]. As such, neural networks need to be tightly coordinated. Glia-mediated control of axonal conduction velocity through myelination is thus of upmost importance because optimal dynamics must be maintained to preserve brain function during development and learning.

Activity-dependent myelin remodelling, through neuron-glia feedback, dynamically coordinates neuron signaling by optimizing conduction velocity and the temporal synchrony of action potentials. But how does activity-dependent myelination optimize synchrony amongst neurons in spite of variability in axon length [5, 22]? From a network perspective, conduction delays between neurons/populations should be roughly equal with each other to guarantee synchrony (i.e., temporally coincident action potentials). This implicitly means that conduction velocity should scale with distance and/or axon length (i.e., longer axons should be more myelinated – see Fig. 1), minimizing the variance between conduction delays and aligning action potential timing. To test this in silico, we combined computational and mathematical approaches and developed a deceptively simple recurrent spiking neural network model endowed with neuron-glia feedback.

**Figure 1.**
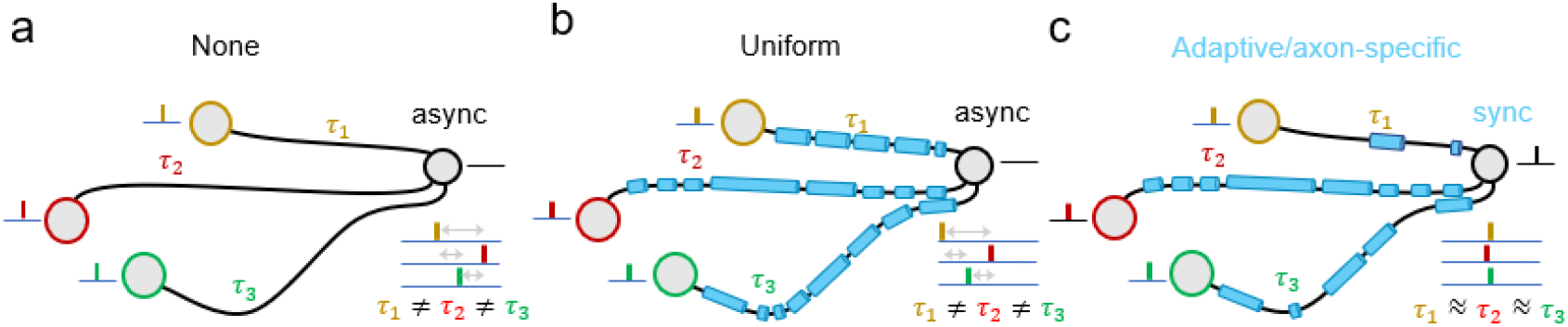
Myelination, axonal length and synchrony. In this illustrative example, pre-synaptic neurons (yellow, red and green circles) are connected with a post-synaptic target cell (black circle) with axons of various lengths. **a**, In absence of myelination, the slow conduction velocity results in conduction delays (*τ*_1_, *τ*_2_ and *τ*_3_) which are proportional to axonal lengths. These delays are dispersed enough to prevent the temporal alignment of propagating action potentials, resulting in asynchronous activation of the post-synaptic neuron. **b**, If myelination is uniform across axons (blue cylinders), the conduction velocity is higher (but finite) and remains proportional to axonal lengths. Delays may remain too variable and action potentials are asynchronous. **c**, If myelination is adaptive and axon-specific, each axon possesses a conduction velocity that scales with its length, yielding similar delays. This results in synchronous action potentials and successful activation of the post-synaptic neuron.

To characterize myelin remodelling, we created a learning rule inspired by experimental data called the Activity-Dependent Myelination (ADM) learning rule. This learning rule is: a)*activity-dependent*: fluctuations in neural firing rates positively regulate myelination; b) *phenomenological* [24]: the net conduction velocity along an axon is assumed to reflect the degree of myelination along that axon; and c) *unsupervised*: no a priori knowledge about optimal myelination structure is used to guide learning. As a first approximation to experimental observations about activity-dependent myelination, the learning rule assumes that conduction velocity scales linearly with firing rates (see Methods).

New experimental findings indicate that, just like synaptic plasticity, myelination is activity-dependent and is influenced by neural activity [6–13]. Optogenetic and electrical manipulations have shown that increases in neuronal firing promote myelin formation and stabilization [6, 7], revealing that neuron-glia interactions play an important yet understudied role in brain plasticity. The plasticity of white matter has been shown to have measurable physiological and behavioral impact in both animal models [8–10], and human studies [14–16]. For example, in rodent studies, it has been shown that learning induces de novo myelination in motor circuits [8, 17], and circuits underlying spatial memory [9, 10]. In humans, it has been shown that activity-dependent myelination plays a key role in homeostasis [14, 15]. Computational work based on human data has also shown that activity-dependent myelination can preserve oscillatory activity in the presence of disease [18]. Identifying the neurophysiological mechanisms involved in neuron-glia interactions and activity-dependent myelination is a focus of intense research [6, 7, 11]. Experiments collectively point towards a relationship between myelin remodelling and neuronal firing, where axonal conduction velocity (and hence delay) may change bidirectionally in a use-dependent way [6, 12]. Studies have shown increased firing rates result in increased myelination, and decreased firing results in myelin retraction [6, 19–21].

Building on these findings to generalize the results to spiking networks, we equipped every axonal connection of our network model with conduction velocity obeying the ADM rule and explored how it shaped temporal synchrony between neurons. We did this to: (1) gain insight about the role played by neuron-glia feedback and activity-dependent myelination on synchrony; (2) disambiguate the respective contributions of synaptic versus axonal/glial plasticity with respect to synchrony and homeostasis. Synaptic connectivity in the model was tuned so that, in the absence of time delays (i.e., zero distance between neurons or infinite conduction velocity), the network exhibits synchrony: firing rates undergo correlated and synchronous jumps between quiescent and active states, mimicking fluctuations observed in vivo [25, 26]. We then embedded this network in a three-dimensional volume and assigned uniform conduction velocities for all network connections, exposing axonal connections to conduction delays of various durations – mirroring variability in axon length. Unsurprisingly, resulting delays suppressed correlated activity and pushed the network away from the synchronous state, despite strong synaptic coupling. Introducing plasticity through the ADM rule, network dynamics homeostatically converged back towards synchrony.

Furthermore, we tested the influence of the ADM rule on network synchrony under two physiologically relevant conditions: when the network geometry changes to emulate development or disease, and as the network is perturbed by external stimuli, to model experience-dependent circuit modifications (or learning). We found that the ADM rule made the synchronous regime scalable: synchronous dynamics were robustly preserved despite changes in network volumes that scaled over four orders of magnitude. Most importantly, changes in conduction velocities observed during learning were not uniformly distributed across network connections: modifications in conduction velocities were instead found to be axon-specific and length-dependent. Second, exposing the network to external stimuli (excitatory or inhibitory) to mimic inputs resulting from experience or learning, our simulations show that axonal conduction velocities and delays could be bidirectionally and reversibly tuned through the ADM rule.

## 2 Results

### Variability in Conduction Delays Suppresses Synchrony

To model the effect of axon-glia feedback and myelin remodelling, we devised a learning rule, referred thereafter as the activity-dependent myelination rule (ADM; see Methods for details), in which conduction velocity (*c_ij_*) along the axonal connection between neuron *i* and neuron *j* changes as a function of the firing rates passing along that axon (see Fig. 2a). This rule combines two elements: i) a positive, firing-rate dependent term (*f_ij_*) that increases the conduction velocity as activity passing along an axon increases – representing myelin formation; and ii) a negative decay term (*r_ij_*), portraying the decrease of conduction velocity in absence of neural activity – representing myelin retraction. Combined, these two components compete at the level of individual axons, resulting in a dynamic and adaptive feedback between neural firing rate and conduction velocities.

**Figure 2.**
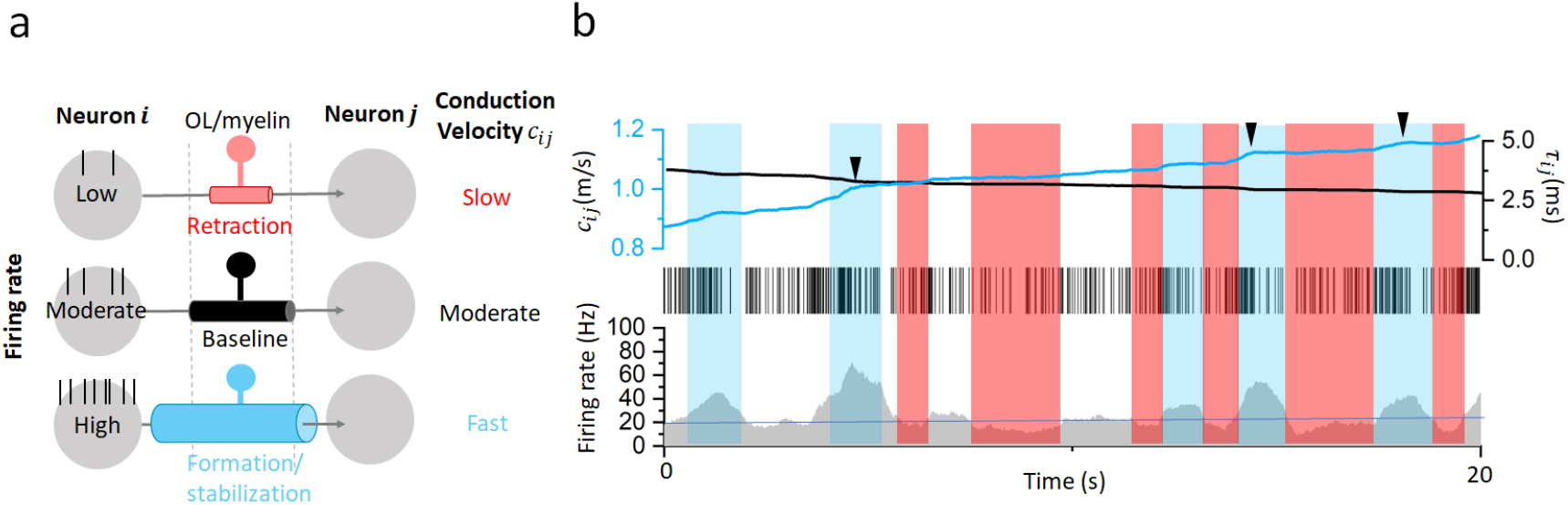
Representative dynamics in conduction velocity resulting from the ADM rule. **a**, Sketch of the ADM rule. Neuron *j* receives signals from neuron *i* (both neurons are depicted as grey circles) with different intensity. Myelin retraction (red cylinder) occurs at low firing rates resulting in slow conduction velocity, while myelin formation (blue cylinder) occurs at high firing rates resulting in fast conduction velocity. **b**, Firing rate fluctuations across an axon result in jumps in net conduction velocity which, collectively, change axonal conduction delays. Sustained spiking promotes myelin formation (blue rectangles), while periods of silence promote myelin retraction (red rectangles). Because formation and retraction occur at different rates, their magnitude is not equal and the overall conduction speed increases until it stabilizes. Parameters used are *α_retraction_* = 0.0001 *ms*, *α_formation_* = 0.001 *ms*, Ω = 10 *mm*, *w_o_* = 0.21, *h* = 0.1, *β* = 25, *ρ* = 0.15, *c_min_* = 0.1 *m/s* and *c_max_* = 100 *m/s*. The same values of the parameters are used throughout, unless otherwise noted.

We set all conduction velocities to the same baseline (before learning) uniform initial value of *c_o_* = 0.1*m/s* (for all axons). This baseline conduction velocity sits inside the range of unmyelinated and/or minimally myelinated axons [27, 28].

We tuned synaptic connectivity in our model such that when distances between neurons are zero (i.e., zero conduction delay and no influence of conduction velocity), the network resides in a synchronous regime. The appropriate values of synaptic gain and connection probability parameters were determined mathematically. For such dense synaptic coupling, synchrony (i.e., temporal alignment of action potential fluctuations) is expected due to a phenomenon called multistability. Multistability leads to noise-induced jumps in firing rates which flip between quiescent and active states (see Methods). Such dynamics notably exhibit critical-like features, reminiscent of those observed in vivo [25, 26].

Does synchrony, resulting from dense synaptic connectivity, persist in the presence of conduction delays? To explore this, we introduced variability in axonal lengths to the model by spatially distributing the neurons randomly in a cubic volume of Ω^3^ = 10^3^ *mm*^3^ (here Ω denotes the edge of a cube). Axonal lengths, resulting from the physical distance between the neurons, were approximated by the Euclidian distance. Introducing such non-zero distances between the neurons resulted in axonal conduction delays. Those delays (*τ*) were computed as the ratio of the Euclidian distance (*l*) between neurons over the conduction velocity (*c*) for that particular axonal connection, i.e., *τ* = *l/c*. Through this process, we introduced significant variability in conduction delays and length-dependent jitter in the timing of action potentials (see Fig. 1). We note that throughout, synaptic gains and connection probability remained unchanged.

We first performed numerical simulations of our spatially distributed baseline model network in the absence of the ADM rule – to serve as a control condition. As shown in Fig. 3a, without activity-dependent changes in conduction velocity, network activity was found to be asynchronous and characterized by low firing rates. Despite dense synaptic connectivity, the uniform and slow conduction velocity (*c_o_*) of this baseline model resulted in significant variability in conduction delay, which suppressed synchrony, as described above. This result was expected from the previous theoretical studies on delayed networks [2, 29]. This confirms that conduction delay variability, resulting from inadequate conduction velocity distribution, represents a major obstacle to coincident neural signaling and synchrony, despite strong synaptic coupling.

**Figure 3.**
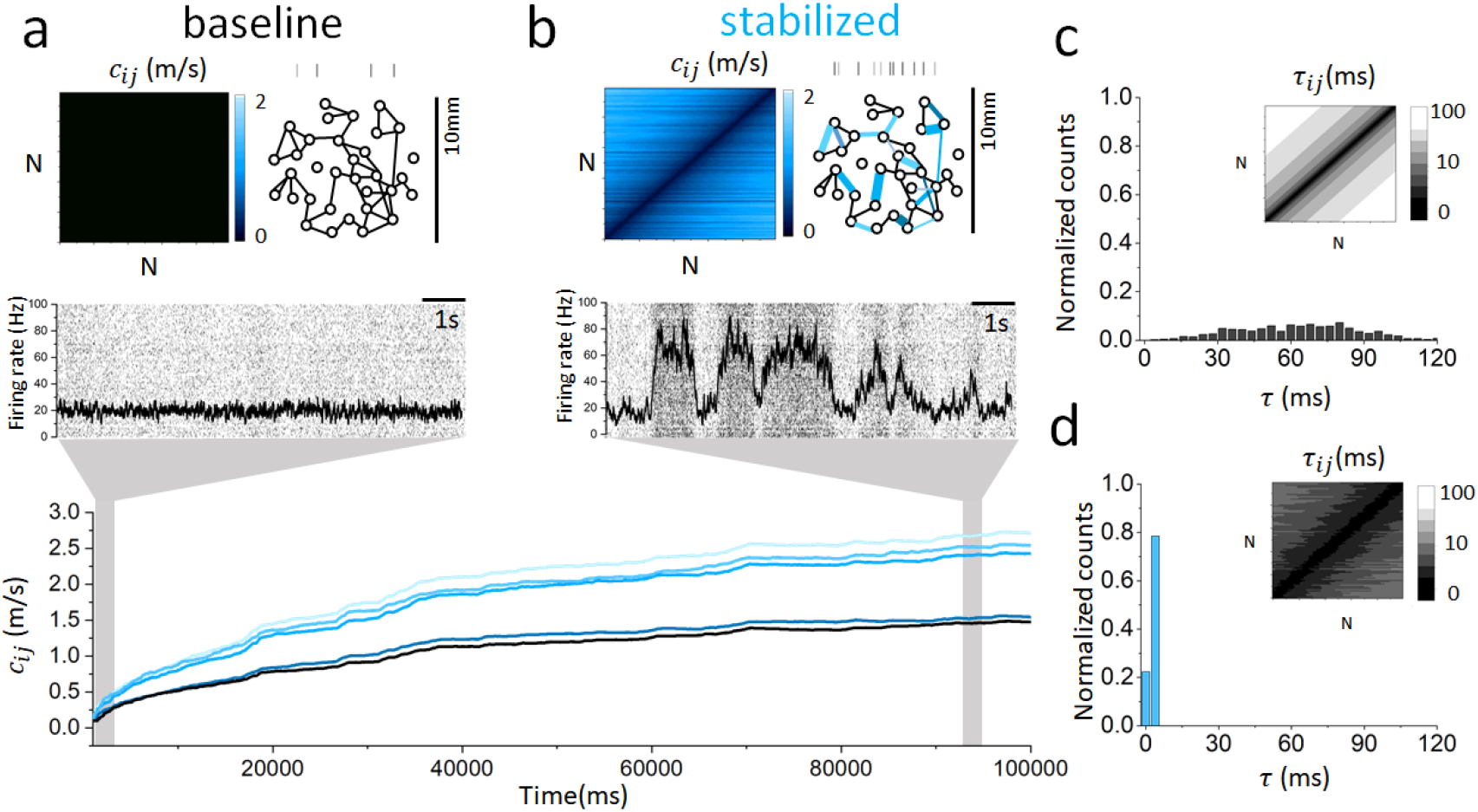
Activity-dependent myelination (ADM) tunes the network towards synchrony. **a**, Network axonal connections were initialized with baseline conduction velocity of *co* = 0.1*m/s*. Neurons were distributed randomly in a cubic volume with dimensions Ω^3^ = 10^3^ *mm*^3^. The resulting dynamics are characterized by asynchronous and low firing rates. **b**, After a period of 10^5^ *ms* adapting under the action of the ADM rule, the conduction velocity increased and plateaued, leading the network dynamics towards synchrony. **c**, Without the ADM rule, baseline conduction delays were highly variable and conditioned by the axonal tract lengths set up randomly by the Euclidean distance between neurons. **d**, After ADM-based learning, the conduction delays are both less variable and statistically shorter, as a direct consequence of increased conduction velocities.

### Homeostatic Control on Synchrony

We then tested whether the introduction of the ADM rule to our baseline network model would restore synchrony. How can adaptive and activity-dependent changes in conduction velocity autonomously restore synchrony in the network? Can it be used to compensate for variability in axonal lengths as physical distances change such as during development? To answer these questions, we let the baseline system evolve and adapt as we monitored its activity. As seen in Fig. 3b, after a simulation period of 10^5^*ms*, the system converged back to synchrony autonomously. Conduction velocities throughout the network stabilized: they first increased and then plateaued. As a direct consequence, both the mean and variance of the conduction delay distribution decreased (Fig. 3c and 3d). The results depicted in Fig. 3 demonstrate that adaptive changes in conduction velocity amplify firing rates and foster synchrony.

### Adaptive Myelination Compensates for Changes in Network Geometry

One of the important roles of myelination is to compensate for increasing physical distances in the developing brain. We tested whether the results of Fig. 3 could be generalized across different spatial scales, and whether ADM-based learning preserved synchrony despite changes in network geometry. By geometry, we refer to changes in the network spatial scale and axonal lengths.

We thus repeated the numerical experiments in Fig. 3 and let the system stabilize, but on geometries – or spatial scales – that spanned four orders of magnitude (from Ω = 1 *mm* to Ω = 10^3^ *mm*). Importantly, synaptic connectivity remained unchanged throughout – allowing us to dissociate the role of synaptic versus glial plasticity in maintaining synchrony.

Our simulations demonstrated that the network autonomously adjusted its conduction velocity statistics to compensate for increased spatial distances. Indeed, as can be seen in Fig. 4, synchrony was achieved irrespective of the spatial scale considered. The ADM rule tuned the conduction velocities until synchronous correlated fluctuations in firing rate emerged. Despite increasing variability in axonal lengths, stabilized delays were found to be statistically similar. To quantify this and measure the level of synchrony in the network, we measured the firing rate variance (*σ*^2^) across independent trials and after the same learning period (see Methods). Across the four spatial scales considered, the variance *σ*^2^ remained stable and decreased dramatically only when ADM learning was turned off by setting the learning rate to zero (i.e., *ϵ* = 0 in Eq. (3)).

**Figure 4.**
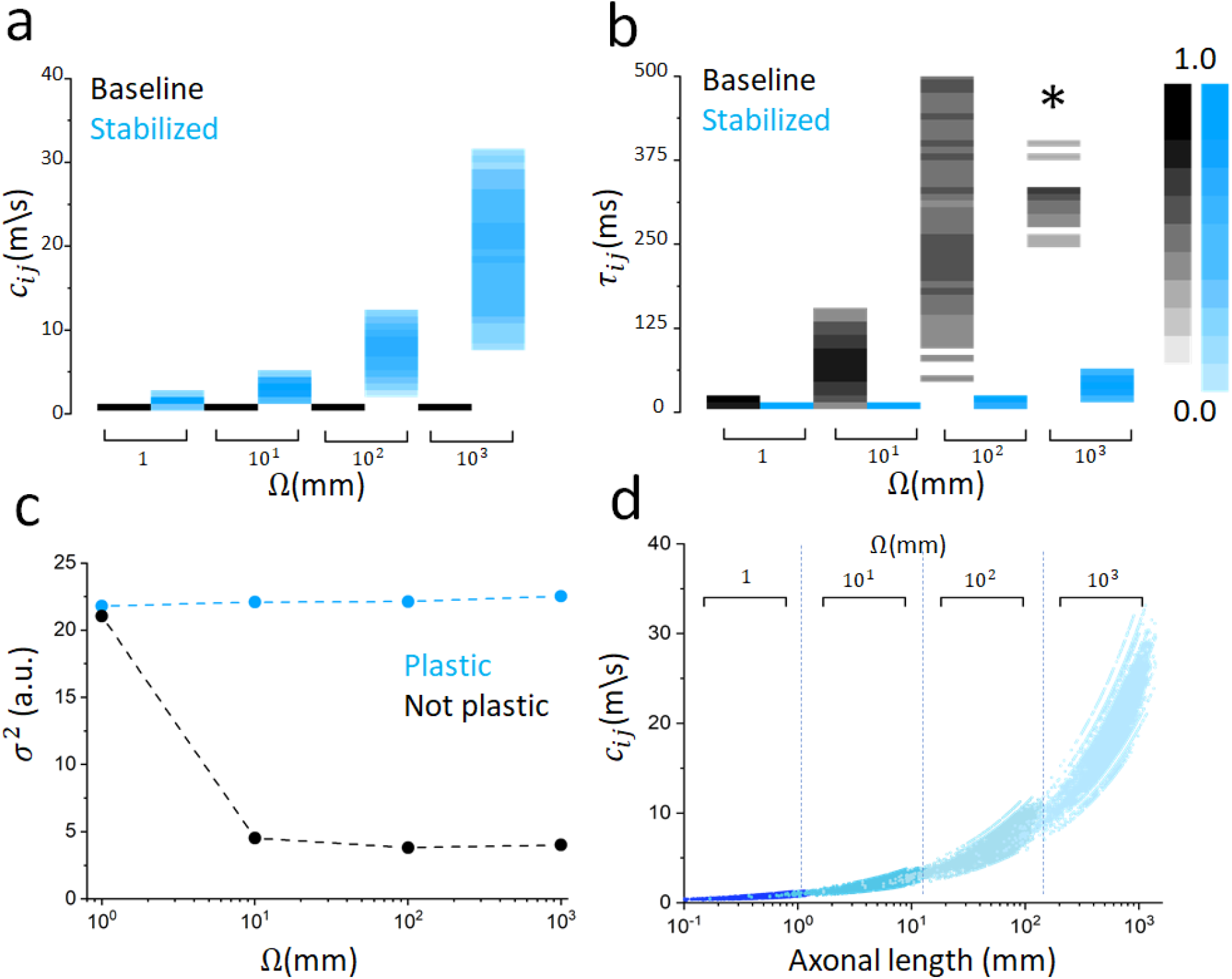
The ADM rule preserves synchrony even as network geometry changes. As the distance between neurons and axonal lengths increase, activity-dependent myelination compensates to maintain synchronous activity. **a**, Conduction velocities scale with network size. As the spatial extent of the network varies across the range Ω = 1 *mm* to Ω = 10^3^ *mm*, the stabilized conduction velocity distribution (after ADM-based learning, blue) shifts towards an increasingly larger range, despite the same initial baseline value (black). **b**, Resulting changes in conduction delays. As the spatial scale increases, the initial baseline distribution is wider (the asterisk denotes that part of the distribution exceeds the range plotted the range examined here and were excluded). This is, however, compensated by adaptive changes in conduction velocity via the ADM rule (Panel a). **c**, Adaptive conduction enables the network to stay in the synchronous state despite changes in physical distances (blue). This scalability property vanishes once ADM rule is turned off (black). Same initial conditions were used throughout the simulations. **d**, Activity-dependent changes in conduction velocity are not uniform, but axon-specific: stabilized conduction velocities increased with axon lengths, across all scales considered in Fig. 3, where Ω ranges from 10° to 10^3^ *mm* (dark to light blue). These axon-specific changes compensate for axon lengths and reduce conduction delay variability.

Our simulations revealed another important consequence of activity-dependent changes in conduction velocity. Despite being uniformly distributed across network connections before adaptation (i.e., baseline), stabilized conduction velocities were not. Indeed, changes in conduction velocities were found to be axon-specific. As can be seen in Fig. 4d, stabilized conduction velocities scaled with axon lengths, across all network scales considered: longer axons became faster compared to shorter ones, compensating for axon length variability and minimizing the variance between conduction delays (cf. Fig. 4ab). These simulations demonstrate that the ADM rule can adapt conduction velocities to preserve synchronous dynamics over a wide range of spatial scales, implementing activity-dependent and axon-specific modifications in conduction velocity.

### Bidirectional and Reversible Influence of Stimuli on Axonal Conduction Velocity

One key experimental observation stemming from multiple recent studies is that stimuli and learning-dependent inputs engage myelination. Optogenetic [6], electromagnetic [7] or learning-dependent [9, 10, 12, 16] stimuli translate into changes in myelin micro-structure. Such myelin remodelling impacts action potential propagation and timing [7]. Changes in myelination have further been shown to be reversible and bidirectional [13, 16], suggesting that stimuli can interfere with neuron-glia feedback loops to influence conduction velocity and hence conduction delays.

We investigated whether stimuli could engage network plasticity in our model to influence axonal conduction velocity. We transiently stimulated neurons in our model and examined how inputs influenced conduction velocities during and after stimulation. We first let the system adapt and stabilize, and applied both excitatory (increasing fire rate) and inhibitory (decreasing fire rate) stimuli of various intensities during a fixed time period. We then measured how conduction velocities and delays changed both during and after stimulation. As can be seen in Fig. 5, prior to stimulation, conduction velocities and associated delays have reached an equilibrium. This equilibrium is perturbed during stimulation. Because the connectivity has stabilized before stimuli onset, network dynamics were synchronous. However, during stimulation, synchrony was replaced by input-driven responses in which firing rates scaled with stimuli amplitudes. Because the ADM rule scales linearly with firing rate, conduction velocities also mirrored stimuli amplitudes. The dynamics observed were reversible: at stimulation offset, conduction velocities converged back to their pre-stimulation values in a homeostatic fashion. This reversibility is reminiscent of myelin retraction observed in absence of stimuli [12, 13, 16]. Firing rates also converged back to their baseline values as stimulation was turned off and synchrony reemerged. Together, these results show that activity-dependent myelination in our model can be bidirectionally and reversibly modulated by stimuli.

**Figure 5.**
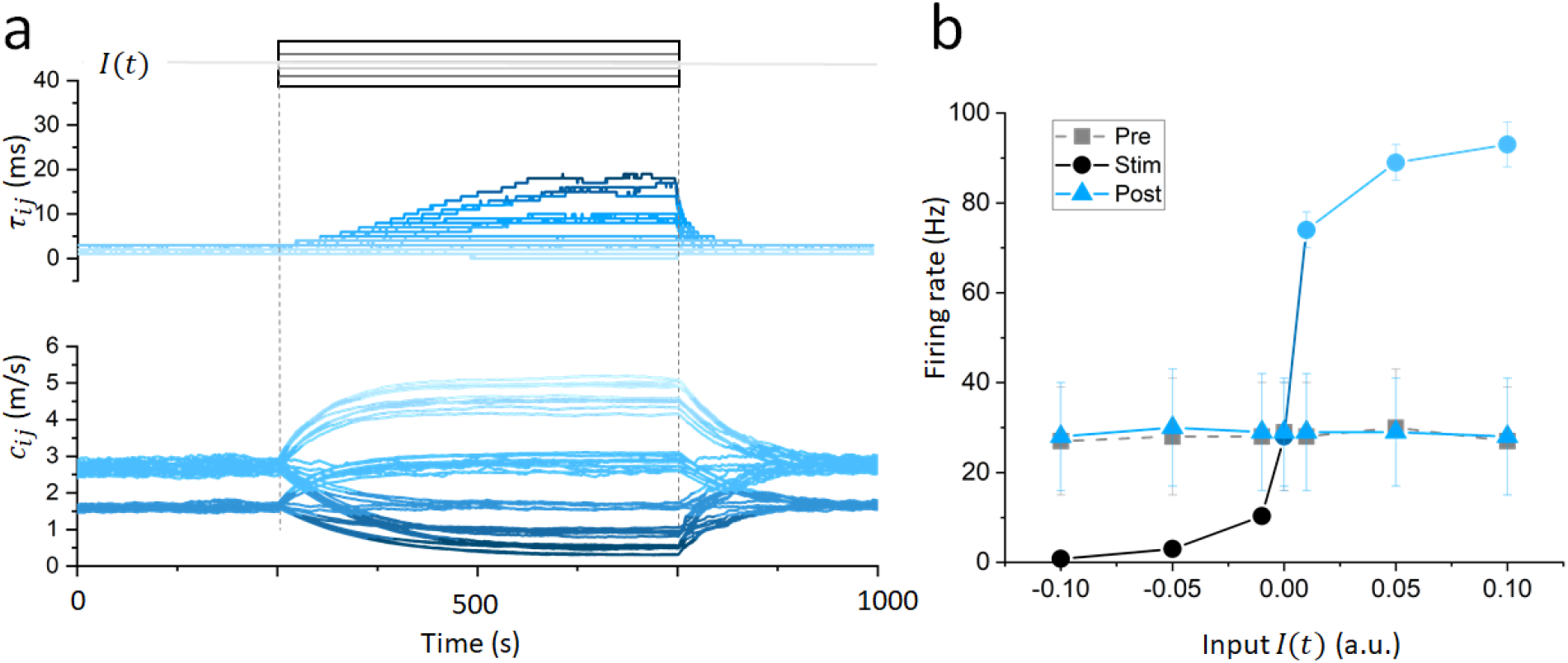
Bidirectional control of conduction velocities and delays by transient stimulation. **a**, As stimuli of various amplitudes (*I* = *−*0.10, *−*0.05, *−*0.01, 0.00, 0.01, 0.05 and 0.10) are delivered uniformly to all neurons in the network, the response triggers adaptive changes in conduction velocities and associated time delays. Increases in activity speed up connections, and the opposite occurs if the firing rate decreases. Because of the inverse relationship between conduction velocities and delays, inhibitory stimuli have a more salient effect on conduction time. As the stimulation is turned off, conductive properties of the various network connections converge back homeostatically to their pre-stimulation values. The blue shading reflects the magnitude of the conduction velocity and the corresponding conduction delays. **b**, Firing rates of the neurons are also tightly regulated. During stimulation, they reflect the stimulation amplitude, but otherwise converge back to their equilibrium values. Network activity was simulated for a total of 1200 *s*, while stimulation was applied during the interval 400 *s* and 800 *s*.

## 3 Discussion

The question that is puzzling especially from the perspective of neural networks, whose structure continuously change due to learning, stimuli and during development, is how neural circuits compensate for variability in axonal lengths to preserve synchronous neural communication. In this work, we hypothesized that neural circuits optimize synchrony through firing rate-dependent changes in axonal conduction velocity. We developed a spiking neural network model enhanced with neuron-glia feedback. To model myelination, we created a learning rule inspired by experimental data and known synaptic plasticity rules [31], called the Activity-Dependent Myelination learning rule (ADM). By this rule, axonal conduction velocity changes linearly with firing rates. We tuned synaptic connections so that network activity is synchronous, and introduced conduction delays to examine their impact on synchrony. Despite strong synaptic coupling, conduction delays induced by variable axonal lengths suppressed synchrony. Once the ADM rule was enabled, our simulations and analysis showed that the network converged back to synchrony autonomously, compensating for axonal length variability. This suggests that synchrony represents a homeostatic target for our network model under the action of the ADM rule.

To generalize these results, we explored the effect of increasing spatial distance between neurons, and altered the geometry of our model network by considering distances ranging over four orders of magnitude. Our results demonstrate that irrespective of the range considered, the ADM rule allowed the network to reach synchrony despite increased variability in axonal lengths and delays. These simulations demonstrated that through ADM learning, synchrony becomes scalable: synchronous activity can be maintained in a homeostatic fashion over a wide range of geometric constraints. Our simulations also showed that despite being uniform before learning, conduction velocity became axon-specific and length-dependent: stabilized conduction velocities scaled with axon lengths. Note that this length dependence of conduction velocity is in line with numerous experimental observations, in which longer axons are usually more myelinated than shorter ones to promote coincident neural signaling (see [32] and references therein).

To examine the influence of inputs resulting from learning or experience, we subjected our network model to both excitatory and inhibitory stimuli of various amplitudes. Our simulations showed that axonal conduction velocities and delays could be bidirectionally and reversibly tuned. At stimuli offset, the conduction velocities converged back to the synchronous state in a homeostatic fashion. This firing rate-dependent change in conduction velocity mirrors the reversible changes in myelination observed experimentally during and after learning [16].

In [18], a large scale brain network model informed by primate structural connectivity data was used to examine how phase-dependent changes in conduction velocity influence global oscillatory activity, notably in presence of injury. Our results extend and generalize these results using an adaptive myelination rule inspired by experimental data [6, 19–21] focused on spiking activity, as opposed to oscillatory responses. While neural oscillations are certainly involved in neural communication, numerous systems do not display neural oscillations while requiring tight temporal coordination (e.g., auditory and electrosensory systems [32]).

Combining these results suggests that activity-dependent myelination endows neural circuit with flexible coordination properties. Because synaptic connectivity was fixed throughout, only the timing of neural interactions changed. As such, it is not the amplitude, but instead the coordination of neural interaction that reinforced synchrony. If we regard synaptic plasticity as a gain control mechanism, activity-dependent myelination represents a timing control property; this speaks of synaptic and axonal/glial plasticity as playing complementary roles in implementing and preserving brain function.

While insightful, our network model and plasticity rule (i.e., the ADM rule) represent trade-offs between biological relevance and mathematical tractability, and we acknowledge the following limitations and opportunities for future study. First, the ADM rule implemented is phenomenological [24] in nature and bypasses the rich neurophysiological mechanisms involved in axon-glia signaling and myelination [6, 13, 33, 34]. The ADM rule was not only inspired by experimental data, but also by decades of work on synaptic plasticity [35, 36]: phenomenological learning rules (e.g., Hebbs learning, STDP) have played a key role in our current understanding of learning and memory. Nonetheless, uncovering the mechanisms involved in neuron-glia feedback remains a topic of intense research. Future experimental work more precisely defining the relationship between neuronal firing and myelination in different brain regions and cell types will help us to refine our model.

Second, we have limited our modelling to the net effect of glia on conduction velocity. Oligodendrocytes, as well as other glial cells (e.g., perinodal astrocytes) are known to significantly impact neural activity [21] and thus likely involved in maintaining homeostasis. This important oversight is left for future work.

Third, the neuron and circuit models used here retain limited abstractions, and do not capture the full richness of action potential emission and propagation. We justify these shortcomings by our desire to assess network-scale (e.g., as opposed to single axon) effects of activity-dependent changes in conduction velocity. Towards this goal, mathematical tractability represents a powerful asset to better understand the contribution of myelination in neural network plasticity.

Fourth, using the Euclidian distance to estimate axonal lengths is commonplace in the computational literature, notably in models informed by connectomic data [1, 3, 4, 18]. It nonetheless remains a coarse approximation. Including detailed anatomy of axonal pathways (e.g., using DTI tractography estimates) would represent a relevant development to improve the link between experiments and modelling.

Fifth, the spatial and temporal scales considered in this study do not necessarily match those involved during and after animal development. Indeed, adaptive myelination occurs over periods of hours, days and years, and thus much slower than what is modelled here [37, 38]. However, the change of time scale does not impact the asymptotic value of the conduction velocities, simply their convergence rate. In addition, the distances considered here certainly go beyond those concerned in many animal neural circuits, but instead represent idealized milestones.

Lastly, our results and conclusions are contingent on the dynamics generated by our model. The synchronous activity observed by our network model results from noise-induced transitions between quiescent and active states. This is one out of many neural activity patterns that could also represent homeostatic targets. The inclusion of both excitatory and inhibitory cells would further enrich the range of possibilities, notably the presence of local and/or global oscillations [18].

## 4 Methods and Materials

### Spiking Network Model

We built a network of *N* recurrently connected excitatory spiking neurons. This microcircuit represents distributed functional network where neurons can be either local (spatially clustered together) or not (distant from one another). This distributed neural assembly is interacting through axonal pathways, which could be thought of as projecting globally through white matter, or locally between cortical layers, for instance. To this end, we uniformly and randomly distributed the neurons location in the cubic volume Ω^3^. The resulting Euclidian distance between neurons was used to determine axonal lengths. Because of spatial distances (i.e., axonal lengths), propagating action potentials experience conduction delays. Given an axonal length between any two neurons, *l*, and a conduction velocity, *c*, the time delay is computed as *τ* = *l/c*.

The spiking response of neurons in the model was modelled using a non-homogeneous Poisson process, where the membrane potential of neuron *i*, *u_i_*(*t*), obeys the following delay differential equation

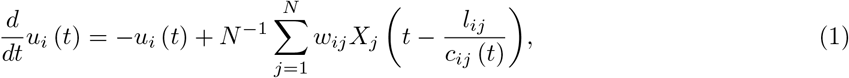

where *X_j_ →* Poisson (*g*[*u_j_*]) is the spike train of neuron *j* with firing rate *g*. The firing rate response function, which maps inputs to firing rate is given here by a sigmoid of the standard form

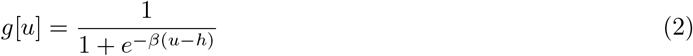

with *β* = 25 and *h* = 0.1. The constants *l_ij_* = *l_ji_* > 0 represent the axonal tract length (i.e., Euclidean distance) between neurons *i* and *j*, and *c_ij_*(*t*) is the conduction velocity. We chose Poisson neurons to account for the intrinsic variability in spike timing in vivo [39].

The synaptic weights were scaled by the connection probability *ρ* = 0.15 and are given by *w_ij_* = *w_o_/ρ*, where *w_o_* = 0.21 is the connection coupling weight. We did this so that the net synaptic connectivity remained invariant to changes in connection probability.

The weights have furthermore been tuned so that the network exhibits synchronous correlated activity in absence of time delays. Mathematically, this means that the weights have been chosen such that the network dynamics sit in a multistable regime and exhibits noise induced jumps between stable steady states, leading to correlated network dynamics as seen in Fig. 3b. While the presence of delays does not impact multistability per se (i.e., steady states of Eq. (1) remain the same with or without delays), it will impact significantly the ability of the network to visit and transition between those states and hence for the network to exhibit synchrony.

### Modeling Neuron-Glia Feedback: the Activity-Dependent Myelination (ADM) Learning Rule

Experimental results show that oligodendrocytes are responsive to changes in neural activity and that myelination is activity-dependent. Optogenetic and electrical manipulations have shown that increases in firing activity promotes myelination [6, 7]. Fluctuations in neural activity engages oligodendroglia, resulting in changes in axonal conduction velocity through the formation of new myelin segments, adaptive changes myelin sheath thickness and/or internodal length [8, 33]. While neuron-glia signaling remain poorly understood and a topic of intense research [6, 7, 11], experiments indicate that neuronal firing rates and axonal conduction velocity are generally positively correlated.

To model such neuron-glia feedback and examine the effect of activity-dependent myelination on neural synchrony, we enhanced our network model in Eq. (1) with an axonal plasticity mechanism. We took a phenomenological approach inspired by Hebbian and use-dependent plasticity [35,36]. Instead of modelling glia directly, we modeled their effect on conduction velocity. We created a phenomenological learning rule – called the Activity-Dependent Myelination rule (ADM). This learning rule is characterized as phenomenological in the sense that along a single axon the conduction velocity reflects myelination.

The ADM learning rule states that the action potential conduction velocity is linearly proportional to the firing rate along a given axon. Specifically, more firing leads to more myelin formation (i.e., conduction velocity increases), and less firing leads to retraction (i.e., conduction velocity decreases). While activity-dependent myelination is most likely not a linear process, the ADM rule is nonetheless a useful simplification for the current model in the absence of more parametric data that speak directly to the relationship between firing rates and myelin remodelling.

Mathematically, the ADM learning rule obeys the following linear ordinary differential equation of the net conduction velocity *c_ij_*:

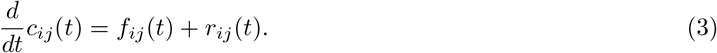

The conduction velocity *c_ij_* along an axonal pathway between neurons *j* and *i* changes according to the interplay of two components: *myelin formation*, *f_ij_*(*t*), and *myelin retraction*, *r_ij_*(*t*).

The myelin formation term is defined as

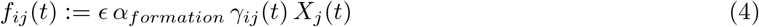

and represents the positive influence of action potentials *X_j_* traveling from neuron *j* to neuron *i* on the conduction velocity *c_ij_*. The coefficient *ϵ* = 0.3 – analogous to the learning rate – sets the gain of how much conduction velocity changes as a function of neural activity. It was chosen so that changes in conduction velocity remain within a relevant physiological range [27, 28]. The rate *α_formation_* = 0.001 *ms* represents a timescale of myelin formation induced by neural firing. Taking into account that the range of potential change in conduction velocity on longer axons is wider than in shorter axons, we added the coefficients *γ_ij_*(*t*) = *l_ij_/c_ij_*(*t*) to compensate for that length-dependent changes in conduction velocity. In addition, *γ_ij_*(*t*) weights the declining impact of increasing conduction velocity with respect to time delays.

The second component of Eq. (3), i.e., the myelin retraction *r_ij_*(*t*), is given by

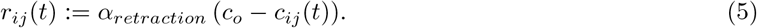

This negative term represents the decrease of conduction velocity in absence of neural activity – identified with activity-dependent myelin retraction. The coefficient *α_retraction_* = 0.0001 *ms* sets the timescale of this decrease. A baseline conduction velocity, *c_o_* = 0.1 *m/s*, was added to model the minimal conduction velocity along an axon. The term *r_ij_* was included in the ADM rule to reflect metabolic demands and resulting myelin retraction in absence of neural activity, in which the conduction velocity decreases exponentially towards a baseline velocity *c_o_* with a rate *α_retraction_*.

Combined, the myelin formation and myelin retraction terms set an equilibrium conduction velocity that is specific to one particular network axonal connection and mirror the neural activity passing through it. Notice that the ADM rule, Eq. (3), models processes that occur at much slower timescales than the activity of the individual neurons. This can be seen from the values of the parameters *α_formation_ ≪* 1 and *α_retraction_ ≪* 1.

According to the ADM learning rule, and as described above, single action potentials *X_j_*, have a positive influence on myelin formation and reflected as an (albeit small) increase in conduction velocity. These two components (myelin formation and myelin retraction) shape the conduction velocities and, by corollary, the conduction delays in the network. Specifically, conduction delays, denoted by *τ_ij_*, between any two neurons *i* and *j* were computed by

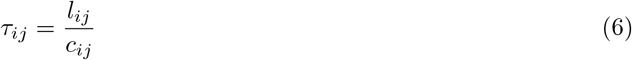

where, recall, *l_ij_* corresponds to the Euclidean distance between these neurons, which have been uniformly and randomly distributed in a cubic volume of size Ω^3^, and *c_ij_* is the conduction velocity determined as per Eq. (3). The influence of spiking activity on the conduction time is immediate and the effective conduction delay for this connection decreases. In absence of spiking, the opposite occurs: conduction velocity stagnates and/or decreases and time delay increases. As can be seen in Fig. 2b, baseline spiking, even in absence of synchrony, increases the conduction velocity. However, more significant increases in conduction velocity occur during correlated and/or synchronous fluctuations.

Combining the ADM learning rule with the network model of Eq. (1) implements a feedback loop in which neural activity influences axonal conduction, and vice-versa, and allows us to examine its impact on network synchrony.

### Quantifying the Variance in Network Mean Activity

To quantify the network response and compare its dynamics with and without activity-dependent changes in conduction velocity, we computed the variance of the mean firing rate. Specifically, we computed

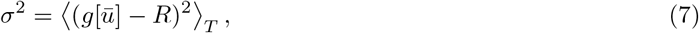

where 〈·〉*_T_* represents the average over trial time *T*, 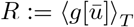 and

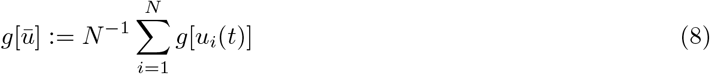

represents the ensemble average response. The variance *σ*^2^ of Eq. (7) is depicted in Fig. 4c.

### Conduction Velocity, Synchrony and the Effect of Time Delays

Our prime focus is to understand the multi scale properties of adaptive myelination. Hence, we are not directly modelling the specific biophysical mechanisms involved in axon-glia feedback [6,13,40] or oligodendrogenesis [9]. While myelination patterns observed in experiments are variable and diverse [23], we choose to focus on the net resulting conductive properties of axons. The conduction velocity over a given axonal tract of length L can vary among other factors as a function of myelin thickness (*t*), segment length (*l*) and nodes of Ranvier density (*R*). Here we consider the net conduction velocity given by

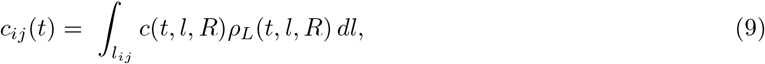

where *ρ_L_* is the probability distribution of those different parameters over the axonal length. The above equation further implies that multiple myelin distribution patterns can lead to identical conduction speeds.

In our model, the homeostatic target state corresponds to synchrony: neural firing rates fluctuate in a collectively correlated fashion, no a priori periodicity is observed. The mean network activity undergoes jumps between quiescent and active states, driven by noise and due to multistability. The presence of time delays influences the probability for such transitions to occur.

To better understand how such fluctuations occur in the network and how they relate to conduction velocity, we may use a mean field approach and consider the collective average behavior of Eq. (1). Assuming only *K ≈ ρN* connections are non-zero, we may rewrite Eq. (1) as

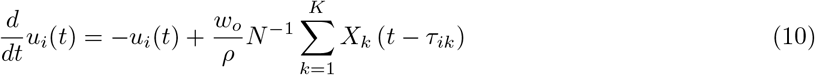

for some 0 < *K < N*. Recall that the delays *τ_ik_* are defined as in Eq. (6), *ρ* is the connection probability and the spike trains *X_k_ →* Poisson (*g*[*u_k_*]).

If we assume that the spike trains are uncorrelated, the sum on the right hand side of Eq. (10) corresponds to a composite non-homogeneous Poisson process, defined by 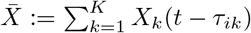, for which the rate is equal to the sum of the individual firing rates (*X_k_*), that is, 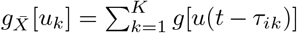. For sufficiently high firing rates, 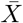 can be approximated by a Gaussian white noise process with mean and variance 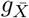,

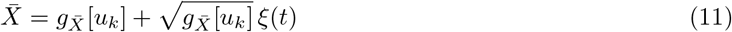

where *ξ*(*t*) is a Gaussian white noise process such that 〈*ξ*(*s*) *ξ*(*s′*)〉 = *δ*(*s − s′*).

In a mean-driven regime, it is possible to write the local membrane potential *u_k_* as a small deviation from the ensemble average: 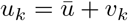, where 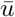 is the ensemble average membrane potential defined as in Eq. (8), and *v_k_* is a small deviation from the mean. Applying this decomposition into Eq. (11), expanding the resulting equation and assuming that 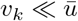 we obtain

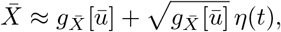

where *η*(*t*) is a Gaussian white noise process such that 〈*η*(*s*) *η*(*s′*)〉 = *δ*(*s − s′*). In the derivation above, higher order terms can be neglected whenever *β* is small enough – which occurs whenever the response function *g* is smooth [41, 42].

Let us now consider

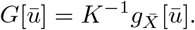

Given that the above calculation holds for all *i* and that *K ≈ ρN*, we receive the following nonlinear stochastic delay differential equation with finite multiplicative noise

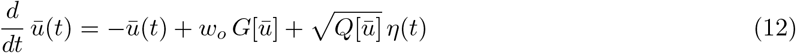

where 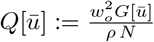. Taking the limit as *N* → ∞, as the number of neurons *N* in the network becomes very large, Eq. (12) is simplified to

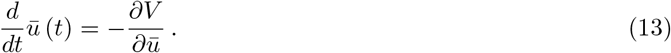

Here 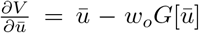, and 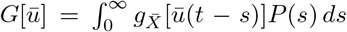, with *P* (*s*) the probability distribution of conduction delays in the network. Eq. (13) is a noise-free equation for the mean activity.

The dynamics observed in the network are analogous to those of a particle randomly fluctuating in a potential well. Indeed, Eq. (13) specifies that the network mean dynamics evolve in a potential *V* with optima given by the steady states, *ϕ^μ^, μ* = 1, 2, 3. The system possesses three steady states due to the choice of the parameters of *g*, as shown in Fig. 6a. In other words, the potential *V* is cubic.

**Figure 6.**
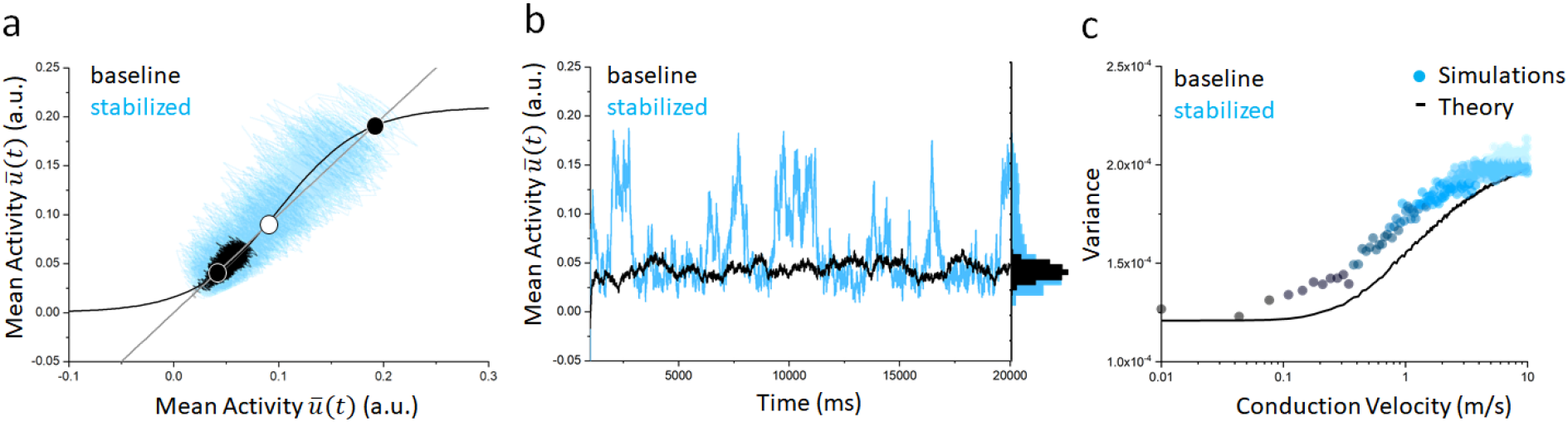
Effect of time delays on mean network dynamics. **a**, Fixed points of the network mean activity. Stable (resp. unstable) steady states are labelled as black (resp. white) circles. **b**, Representative dynamics for baseline (black) and stabilized (blue) conduction velocities. Large conduction delays lead to small amplitude activity that does not permit correlated synchronous fluctuations. The network mean activity remains close to the low activity equilibrium state. Whenever conduction velocities increase due to the ADM rule, mean activity fluctuations increase, and the network undergoes noise-induced jumps between the stable states, resulting in synchrony. **c**, Effect of conduction velocity on mean activity variance. As the conduction velocity increases, conduction delays decrease, amplifying mean activity fluctuations: the variance increases. Simulation data resulting from linearization around the stable steady state plotted alongside theoretical predictions.

The three steady states are solutions to the equation

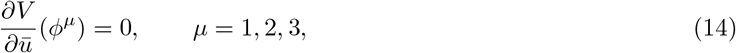

or equivalently, *ϕ^μ^ − w_o_G*[*ϕ^μ^*] = 0. The points *ϕ*^1^ and *ϕ*^3^ are stable, while *ϕ*^2^ is unstable. The first stable steady state *ϕ*^1^ corresponds to a state of quiescence and thus low firing rates. The second stable steady state *ϕ*^3^ corresponds to a state of active neural firing and thus elevated firing rates. These are quiescent and active states visited in Fig. 3b. The fixed point *ϕ*^2^ (white circle in Fig. 6a) is unstable, and delineates the basins of attraction of the two stable steady states.

Our main goal is to understand the role of the delays to transitions between different states leading to synchronization. In [43], the author uses a Fokker-Planck approach to derive the stationary probability density for nonlinear stochastic delay differential equations. To observe intuitively the changes to the system caused by delays we will use a linear approximation. Consider small fluctuations, *v*(*t*), around the stable steady state *ϕ*^1^ such that the solution 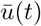 of Eq. (12) takes the form 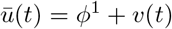. Linearizing Eq. (12) about *ϕ*^1^ yields

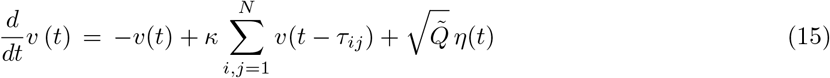

where *κ* ≔ *w_o_ N ^−^*^2^*g′*(*ϕ*^1^), and 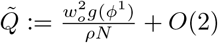. Since *κ* = *O*(*δ*) with *δ ≪* 1, the perturbation approach of [44] is applicable. Therefore, the variance of the dynamics of Eq. (15) is

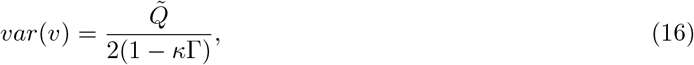

where 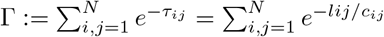.

Fig. 6c depicts the simulated variance of Eq. (15) and the variance in Eq. (16). Observe that the variance is proportional to conduction velocities, or equivalently is inversely proportional to the time delays. That is, long conduction delays (i.e., prior to ADM induced changes when conduction velocities are small) will result in low variance. Fluctuations around the stable steady states will have small amplitude and no network-wide correlated fluctuations (i.e., synchrony) occur. Network firing rates remain low and there is no synchrony. This is the dynamics portrayed in Fig. 3a. In contrast, when conduction delays are small (i.e., after stabilization once velocities are high), the opposite occurs: synchronous fluctuations take place as the mean activity jumps between the stable steady states. This is what is portrayed in Fig. 3b. Representative dynamics for these two cases are plotted in Fig. 6b.

**Table 1.** List of model parameters.

| Symbol | Definition | Value |
| --- | --- | --- |
| $N$ | network size | 100 |
| $\Omega^3$ | cubic volume | $10^3 \text{ mm}^3$ |
| $\epsilon$ | learning rate | 0.3 |
| $\alpha_{retraction}$ | myelin retraction rate | $0.0001 \text{ ms}$ |
| $\alpha_{formation}$ | myelin formation rate | $0.001 \text{ ms}$ |
| $h$ | the $u$ value of the sigmoid's midpoint | 0.1 |
| $\beta$ | steepness of the sigmoid function | 25 |
| $w_o$ | connection coupling weight | 0.21 |
| $\rho$ | connection probability | 0.15 |
| $c_{min}$ | minimal conduction velocity | $0.1 \text{ m/s}$ |
| $c_{max}$ | maximal conduction velocity | $100 \text{ m/s}$ |
| $l_{ij}$ | axonal length | variable |
| $c_{ij}$ | conduction velocity | variable |
| $\tau_{ij}$ | conduction delay | variable |
| $w_{ij}$ | connection weights | variable |
| $c_o$ | baseline conduction velocity | $0.1 \text{ m/s}$ |
| $\gamma_{ij}$ | weight coefficient | variable |
| $\sigma^2$ | variance of the mean activity | a.u. |
| $\text{var}(\mathbf{v})$ | variance of the linear dynamics | a.u. |

## 6 Author Contributions

JL designed the study. JL and AT performed the mathematical derivations and simulations. JL, PF, DM and AT wrote the manuscript.

